# Sulcal variability in anterior lateral prefrontal cortex contributes to variability in reasoning performance among young adults

**DOI:** 10.1101/2023.02.10.528061

**Authors:** Ethan H. Willbrand, Samantha Jackson, Szeshuen Chen, Catherine B. Hathaway, Willa I. Voorhies, Silvia A. Bunge, Kevin S. Weiner

**Author notes:** Co-senior authors (Silvia A. Bunge, Kevin S. Weiner). **Corresponding authors:** Silvia A. Bunge and Kevin S. Weiner **Emails:**.

## Abstract

Identifying structure-function correspondences is a major goal among biologists, cognitive neuroscientists, and brain mappers. Recent studies have identified relationships between performance on cognitive tasks and the presence or absence of small, shallow indentations, or sulci, of the human brain. Building on the previous finding that the presence of one such sulcus in the left anterior lateral prefrontal cortex (aLPFC) was related to reasoning task performance in children and adolescents, we tested whether this relationship extended to a different sample, age group, and reasoning task. As predicted, the presence of this aLPFC sulcus—the ventral para-intermediate frontal sulcus—was also associated with higher reasoning scores in young adults (ages 22-36). These findings have not only direct developmental, but also evolutionary relevance—as recent work shows that the pimfs-v is exceedingly rare in chimpanzees. Thus, the pimfs-v is a novel developmental, cognitive, and evolutionarily relevant feature that should be considered in future studies examining how the complex relationships among multiscale anatomical and functional features of the brain give rise to abstract thought.

## Introduction

Identifying structure-function correspondences is a major goal across subdisciplines in the biological sciences. In neurobiology and cognitive neuroscience, there is broad interest in uncovering relationships between neuroanatomical features of the human brain and cognition— especially for structures in parts of the brain that are largely human-specific. Given that 60-70% of the human cerebral cortex is buried in indentations, or sulci (Zilles et al. 1988, 2013; Van Essen 2007), there is continued interest in the relationships among sulcal morphology, functional representations, and cognition. Previous work exploring this relationship has largely focused on the consistent and prominent sulci within primary sensory cortices (Yousry et al. 1997; Boling et al. 1999; Hinds et al. 2008; Cykowski et al. 2008; Li et al. 2010; Wandell and Winawer 2011; Sun et al. 2012; Benson et al. 2012). Nevertheless, recent work has begun to explore the small, shallow, and more variable sulci in association cortices that are not always present in a given hemisphere. For example, recent studies reveal relationships between the presence or absence of specific sulci in association cortices and individual differences in human cognitive abilities and clinical conditions (for review see Cachia et al. 2021), which could be mediated by differences in white matter architecture in relation to these sulcal features (Van Essen 1997, 2020; White et al. 2010; Zilles et al. 2013).

In the present study, we focus on the anterior lateral prefrontal cortex (aLPFC) given several sequential findings linking the individual variability of an aLPFC sulcus to cognitive performance. Specifically, our recent work implementing a data-driven approach on 12 LPFC sulci (Petrides 2019; Voorhies et al. 2021) showed that the morphology of an aLPFC sulcus (the para-intermediate frontal sulcus, pimfs) was the best predictor of performance on a widely used test of reasoning in a pediatric sample (ages 6-18) (Voorhies et al. 2021). However, since the pimfs is composed of two variably present components (Voorhies et al. 2021; Willbrand et al. 2022c), we then tested whether the presence of specific pimfs components was also related to reasoning scores in a larger pediatric sample. Indeed, we found that the presence of the ventral pimfs component (pimfs-v) in the left hemisphere, specifically, was associated with better reasoning performance (Willbrand et al. 2022c). Here, we tested—and empirically support—the targeted prediction that this pattern of results across the four pimfs components (left and right pimfs-v and pimfs-d) holds across age groups and studies. Further, as in our pediatric sample, the presence or absence of left pimfs-v was linked to scores on a reasoning task but not to a control task that measures lower-level cognitive performance: processing speed. This extension of our previous result in children and adolescents to a separate sample of adults using different cognitive task variants is a notable finding, given a timely discussion among researchers regarding the reliability and generalizability of brain-behavior relationships (Marek et al. 2022; Gratton et al. 2022; Westlin et al. 2023). It is also notable because it converges with studies implicating aLPFC in various tests of reasoning, as discussed below. In addition to these tests of reasoning and processing speed, we had the opportunity in this sample to test whether performance on a test of working memory that requires transitive inference—a different form of reasoning—varies as a function of left pimfs-v presence. We discuss these findings in the context of (i) the role of aLPFC in reasoning, (ii) hypothesized relationships among the presence/absence of sulci, the morphology of sulci, and the efficiency of network communication contributing to performance on cognitive tasks, and (iii) the translational implications of aLPFC sulcal variability.

## Materials and Methods

### Participants

Data for the younger adult human cohort analyzed in the present study were taken from the Human Connectome Project (HCP) database (https://db.humanconnectome.org). Here we used 72 participants (50% female, 22-36 years old, 90% right-handed). These participants have also been used in our previous work in the posterior lateral prefrontal and posteromedial cortices (Miller et al. 2021b; Willbrand et al. 2022b, 2023ba, c). HCP consortium data were previously acquired using protocols approved by the Washington University Institutional Review Board. Informed consent was obtained from all participants.

### Imaging data acquisition

Anatomical T1-weighted (T1-w) MRI scans (0.7 mm voxel resolution) were obtained in native space from the HCP database as well as cortical reconstructions generated through the HCP’s version of the FreeSurfer pipeline (Dale et al. 1999; Fischl et al. 1999; Glasser et al. 2013). All sulcal labeling and anatomical metric quantification was done on the cortical surface reconstructions of each participant.

### Behavioral data

#### Overview

In addition to structural and functional neuroimaging data, the Human Connectome project also collected a wide range of behavioral metrics (motor, cognitive, sensory, and emotional processes) from the NIH toolbox (Barch et al. 2013) that illustrate a set of core functions relevant to understanding the relationships between human behavior and the brain (task details: https://wiki.humanconnectome.org/display/PublicData/HCP-YA+Data+Dictionary-+Updated+for+the+1200+Subject+Release#HCPYADataDictionaryUpdatedforthe1200SubjectRelease-Instrument). 71 of 72 participants in the present project had behavioral scores. Below we describe the three behavioral tests used.

#### Relational reasoning task

The ability to reason about the patterns, or relations, among disparate pieces of information— i.e., relational reasoning—has long been recognized as central to human reasoning and learning (James 1890a, b; Cattell 1943). Tests of relational reasoning assess the ability to integrate and generalize across multiple pieces of information, and help to predict real-world performance in a variety of domains (Alexander 2016). Here, we used scores obtained for each participant on a measure of relational reasoning, the Penn Progressive Matrices Test from the NIH toolbox (Barch et al. 2013). This test is similar to the classic Raven’s Progressive Matrices (Raven 1941), the WISC-IV Matrix Reasoning task (Wechsler 1949) used in our pediatric sample (Willbrand et al. 2022c), and other task variants that are ubiquitous in assessments of what is often termed “fluid intelligence.” In this task, participants must consider how shapes in a 2×2, 3×3, or 1×5 stimulus array are related to one another, for example, an increase, across a row or column, in the number of lines superimposed on a circle. Specifically, participants must identify the abstract relations among items in the array (Carpenter et al. 1990) and select, among five options, the shape that completes the matrix. The task is composed of 24 different matrices, presented in order of increasing difficulty. Testing is discontinued after five incorrect choices in a row, and the total score is calculated as the number of correct responses.

#### Processing speed task

Participants also completed the Pattern Comparison Processing Speed Test from the NIH toolbox (Barch et al. 2013). This test was designed to measure the speed of cognitive processing based on the participant’s ability to discern whether two adjacent pictures are identical as quickly as possible. Here, participants consider several possible differences (addition/removal of an element or the color or number of elements on the pictures). A yes-no button press is used to determine whether the two stimuli are identical, and the final score corresponds to the number of trials answered correctly during a 90-second period.

#### List sorting working memory test

Participants also completed the List Sorting Working Memory Test from the NIH toolbox (Barch et al. 2013). In this task, each participant sequences different visually and orally presented stimuli (alongside a sound clip and written text for the name of the item) in two conditions: 1-List and 2-List. In the former, participants order a series of objects (food or animals) from smallest to largest. In the latter, participants are presented with both object groups (food *and* animals) and must report the food in order of relative size in real life and then the animals in order of size. Notably, completing this task not only requires working memory manipulation and maintenance but also relational reasoning: to report the items in order of size it is necessary to compare pairs of stimuli and then engage in transitive inference across multiple items (Halford et al. 1998).

### Morphological data

#### Defining the presence and size of the para-intermediate middle frontal sulcus

Individuals typically have anywhere from three to five tertiary sulci within the middle frontal gyrus (MFG) in LPFC (Miller et al. 2021a, b; Voorhies et al. 2021; Yao et al. 2022). The posterior MFG contains three of these sulci, which are present in all participants: the anterior (pmfs-a), intermediate (pmfs-i), and posterior (pmfs-p) components of the posterior middle frontal sulcus (pmfs). In contrast, the tertiary sulcus within the anterior MFG, the para-intermediate middle frontal sulcus (pimfs), is variably present. A given hemisphere can have zero, one, or two pimfs components (**Fig. 1**). As described in prior work (Petrides 2013, 2019; Willbrand et al. 2022c), the dorsal and ventral components of the pimfs (pimfs-d and pimfs-v) were generally defined using the following two-fold criterion: i) the sulci ventrolateral to the horizontal and ventral components of the intermediate middle frontal sulcus, respectively, and ii) superior and/or anterior to the mid-anterior portion of the inferior frontal sulcus.

**Figure 1.**
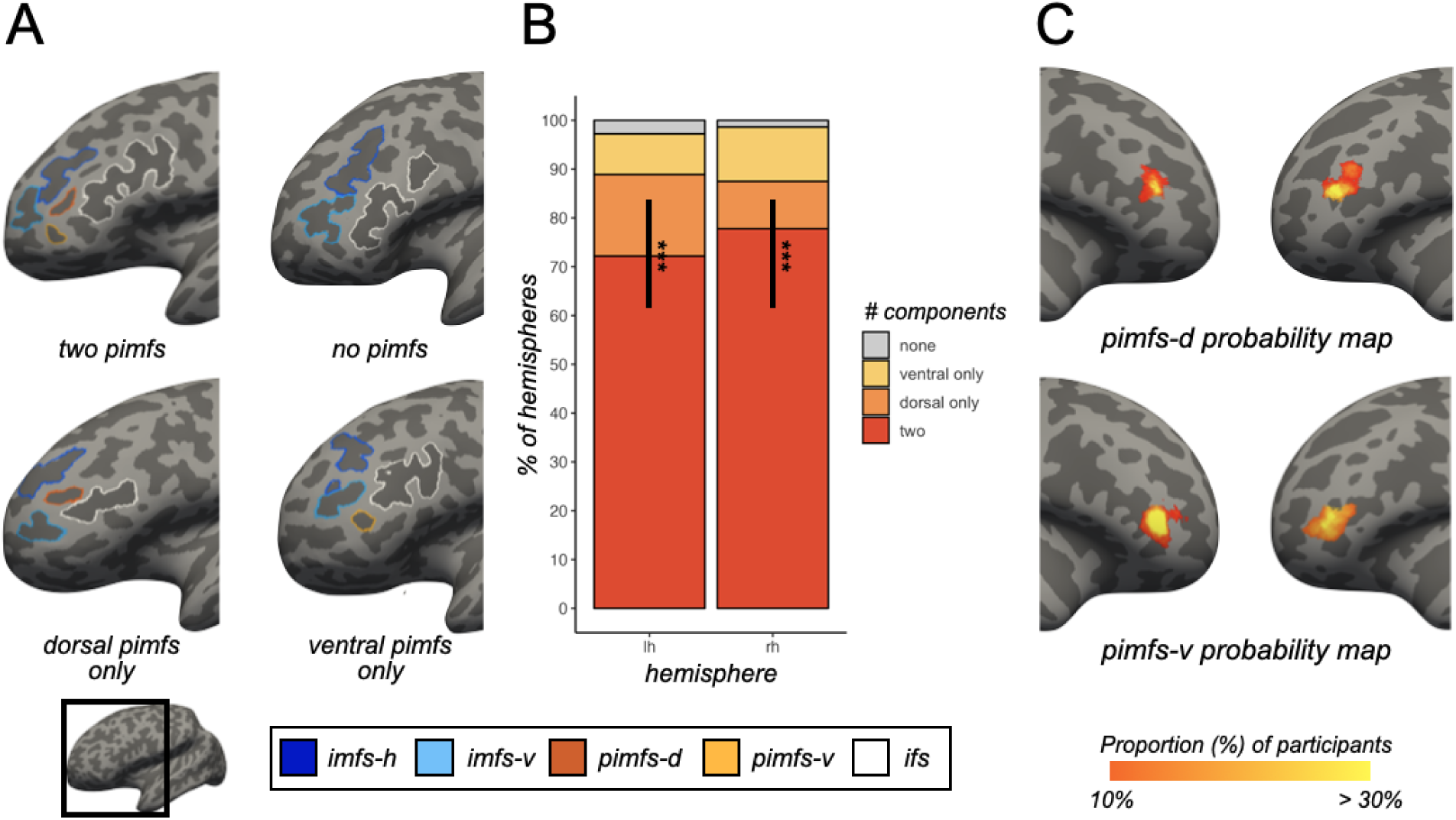
The para-intermediate frontal sulcus (pimfs) is variable across individuals and hemispheres. **A.** Inflated left hemispheres (sulci: dark gray; gyri: light gray; cortical surfaces are not to scale) depicting four types of the pimfs: (i) both pimfs-v/d present, (ii) neither present, (iii) pimfs-d present, (iv) pimfs-v present. Prominent sulci surrounding the pimfs are shown: horizontal (imfs-h) and ventral (imfs-v) intermediate frontal sulci and inferior frontal sulcus (ifs; see legend). **B.** Stacked bar plot depicting pimfs incidence in left (lh) and right (rh) hemispheres (N = 72; see rightward legend). **C.** Maximum probability maps (MPMs) for pimfs-d (top) and pimfs-v (bottom) on the inflated *fsaverage* cortical surface (left surfaces: rh; right surfaces: lh). For visual clarity, the MPMs were thresholded at 10% overlap across participants (the warmer the color, the higher the overlap). (*** *p* < 0.001).

We first manually defined the pimfs within each individual hemisphere with *tksurfer* (Miller et al. 2021b). Manual lines were drawn on the *inflated* cortical surface to define sulci based on the most recent schematics of pimfs and sulcal patterning in LPFC by Petrides (2019), as well as by the *pial* and *smoothwm* surfaces of each individual (Miller et al. 2021b). Using multiple surfaces allows for the establishment of a consensus across surfaces and clearly determines sulcal boundaries. The location of the pimfs was confirmed by three trained independent raters (E.H.W., S.M., S.C.) and finalized by a neuroanatomist (K.S.W.). Although this project focused on a single sulcus, the manual identification of all LPFC sulci (2,877 sulcal definitions across all 144 hemispheres) was required to ensure the most accurate definitions of the pimfs components. For in-depth descriptions of all LPFC sulci, see (Petrides 2019; Miller et al. 2021a, b; Voorhies et al. 2021; Yao et al. 2022; Willbrand et al. 2022a).

The size of the pimfs was quantified as surface area (in mm^2^), as a quantitative comparison to the qualitative metric of sulcal incidence. The surface area of each pimfs label was extracted with the mris_anatomical_stats FreeSurfer function (Fischl and Dale 2000). For participants with two pimfs components, the surface area was quantified as a sum of both components. For participants with one pimfs component, the surface area was quantified as that single component. As in previous work (Clark et al. 2010; Rollins et al. 2020), the surface area was set to zero for participants with no pimfs component. To control for individual differences in brain size, as in prior work (Willbrand et al. 2022c, 2023c; Hathaway et al. 2023), we normalized the surface area as a percent of the hemispheric surface area.

#### Probability map generation

As in prior work (Miller et al. 2020, 2021b; Voorhies et al. 2021; Willbrand et al. 2022b; Hathaway et al. 2022), sulcal probability maps were calculated to display the vertices with the highest alignment across participants for a given sulcus. The label file for each pimfs component was first transformed from the individual to the fsaverage surface. Once transformed to this common template space, the proportion of participants for whom a given vertex is labeled as the given pimfs component was then calculated. For vertices where the pimfs components overlapped, we employed a “winner-take-all” approach. Here, the component with the highest overlap across participants was assigned to that overlapping vertex. With the publication of this article we provide un-thresholded maps and constrained versions of these maps (i.e., maximum probability maps, MPMs) to improve map interpretability. We show the 10% overlap MPMs in **Figure 1C**.

We first compared the incidence rates of the pimfs components—both as number of components (2 vs < 2) and as the presence of the specific pimfs component (dorsal vs ventral)—in each hemisphere with Chi-squared (χ^2^) tests. Note that all statistical tests in the present project were implemented in R (v4.1.2; https://www.r-project.org/) and each set of analyses was corrected for multiple comparisons (with the false-discovery rate; FDR). Chi-squared tests were carried out with the chisq.test function from the stats R package.

### Analysis II: Relating the presence and size of the pimfs to hemisphere, age, and gender

Prior work has indicated that the presence of variable sulci may differ between hemispheres and by participant gender (Paus et al. 1996; Clark et al. 2010; Wei et al. 2017; Amiez et al. 2019, 2021). Further, although sulcal patterning emerges before birth and is stable across the lifespan (Cachia et al. 2021), there could have been an age imbalance in pimfs incidence in our sample, purely by chance. So, too, could there have been an imbalance in pimfs incidence between genders in our sample. Thus, we sought to test for differences in incidence rates as a function of hemisphere, gender, and age. To this end, we implemented three binomial logistic regression GLMs with hemisphere (left, right), age, and gender (male, female), as well as their interactions, as factors for i) number of components [0 (fewer than 2 components), 1 (2 components)], ii) pimfs-d presence [0 (absent), 1 (present)], and iii) pimfs-v presence [0 (absent), 1 (present)]. Finally, in addition to these categorical analyses, we examined a continuous variable, conducting a multiple linear regression for normalized total surface area of the pimfs component(s). GLMs were implemented with the glm function from the stats package. ANOVA χ^2^ tests were applied to each GLM with the Anova function from the car package. Linear regressions were performed with the lm function from the stats package.

### Analysis III: Relating the presence of the pimfs to reasoning performance

These analyses were performed on the 71 participants with behavioral scores. Participant age and gender were not considered in these analyses, as Analysis II showed that they did not differ in terms of pimfs presence (**Results**) or reasoning performance (*age*: ß = −0.01, *p* = 0.98; *gender*: χ^2^ = 0.04, *p* = 0.84). We first ran two-sample t-tests to assess whether the number of components in each hemisphere (*two* vs *less than two*) related to reasoning performance (Penn Progressive Matrices Test). In the main text, we report differences between groups as the percentage increase in average scores (µ) from the “less than two” group to the “two” group with the following equation:

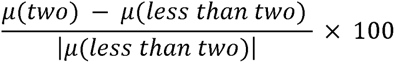

Next, to determine if the presence of a specific pimfs component was related to reasoning performance, we ran additional two-sample t-tests to test for an effect of the presence of the pimfs-v and pimfs-d (*present* vs *absent*) in each hemisphere. As in the prior analysis, we report differences between groups as the percentage increase in µ from the “absent” group to the “present” group.

To determine whether the observed relationship between the left pimfs-v presence and reasoning (**Fig. 2B**) was impacted by differences in sample size between participants with and without this sulcus, we iteratively sampled a subset of participants from the left pimfs-v present group (N = 57) to match that of the left pimfs-v absent group (N = 14) 1,000 times and conducted a Welch’s t-test for each sampling (to account for potential unequal distributions when resampling). To evaluate the bootstrapped effect size, we report the median and 95% confidence interval for the effect size.

**Figure 2.**
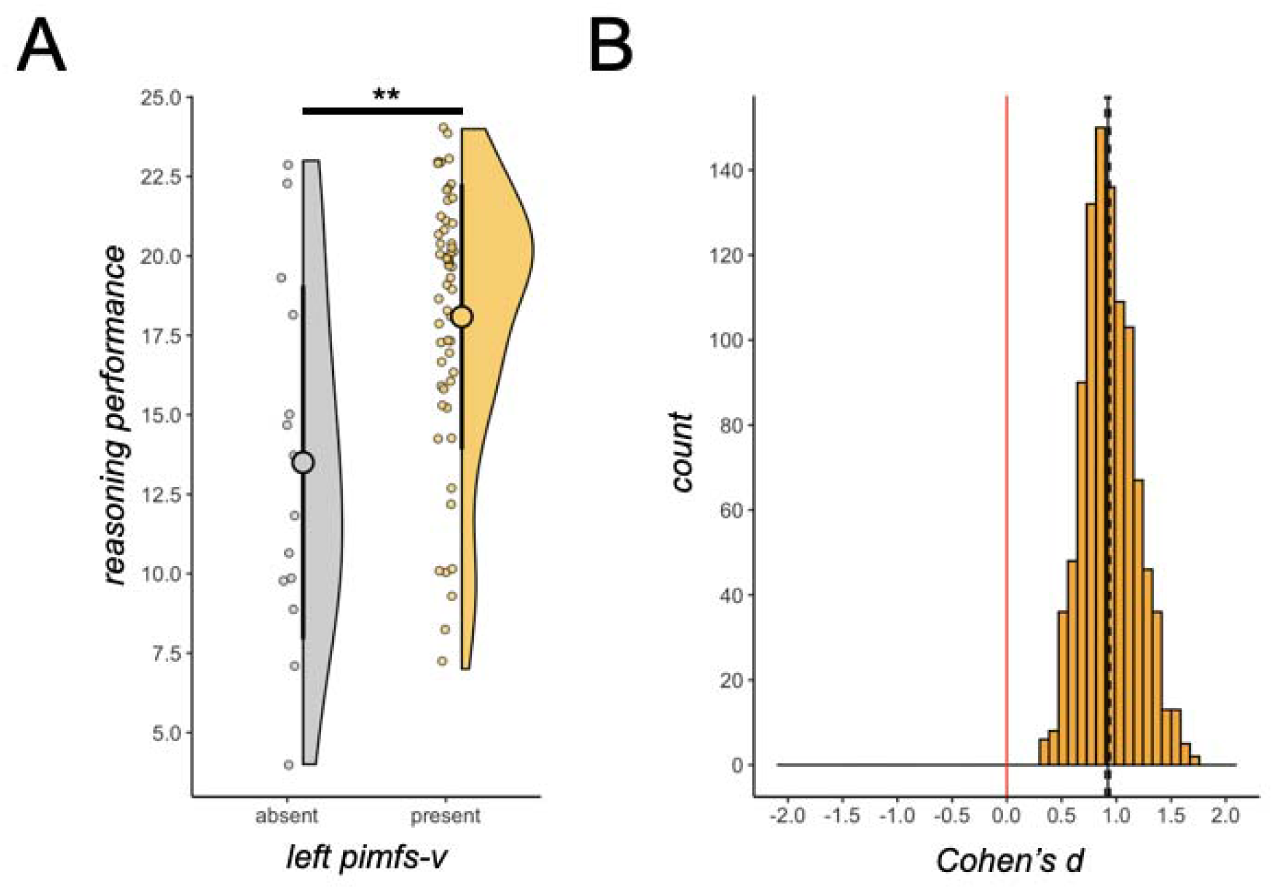
Relational reasoning is related to left pimfs-v presence. **A.** Raincloud plots (Allen et al. 2021) depicting Penn Progressive Matrices task score as a function of left pimfs-v presence in younger adults (present, N = 57; absent, N = 14). Large dots and error bars represent meanlJ±lJstd reasoning score; violin plots represent kernel density estimate. Small dots indicate individual participants. **B.** Histogram visualizing results of the iterative resampling of left pimfs-v presence in A 1,000 times. Distribution of the effect size (Cohen’s d) is shown, along with the median (black line) and 95% CI (dotted lines). Red line corresponds to zero, highlighting that left pimfs-v absence was never associated with higher reasoning scores than left pimfs-v presence. (** *p* < .01).

Next, given the wide age range (22-36 years old) and prior work indicating that reasoning performance peaks in the early 20s (McArdle et al. 2002), we implemented a 3-step procedure to ensure that age did not confound the results and to further evaluate the model fit. First, we ran a linear regression with left pimfs-v presence and age as predictors for reasoning in the full sample (N = 71 with reasoning scores). We then ran a Chi-squared test to compare the previously described regression model to one including age only. Second, to further confirm that age did not drive the effect of left pimfs-v presence on reasoning scores, we conducted variable-ratio matching on age (ratio = 3:1, min = 1, max = 5; optimal ratio parameters were based on the calculation in (Ming and Rosenbaum 2000) with the MatchIt package. Here, the distance between each member of each group (left pimfs-v presence, left pimfs-v absence) was calculated with a logit function:

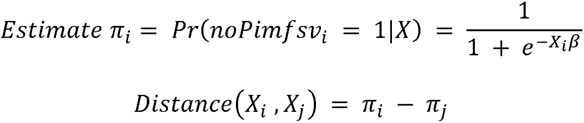

where X is participant age in groups without (i) and with (j) a left pimfs-v. Matches were based on a greedy nearest-neighbor interpolation where each participant in the smaller group (left pimfs-v absent) received 1-5 unique matches from the larger group (left pimfs-v present). Afterwards, a weighted linear regression was run in the matched sample, with left pimfs-v presence and age as predictors of reasoning, to confirm the robustness of our initial finding with the whole sample. Third, we employed a two-pronged analysis to assess and verify the unique variance explained by left pimfs-v presence, while accounting for age-related effects on reasoning. First, we ran a Chi-squared test to compare the previously described weighted-regression model to one with age only. Second, as described and implemented in prior work (Voorhies et al. 2021; Yao et al. 2022; Willbrand et al. 2022c), we fit these weighted-regression models with repeated K-fold cross-validation (CV; five-fold, 10 repeats). Since these are nested models, the best fit was determined as the model with the lowest cross-validated RMSE_cv_ and the highest *R*^2^ value. We did not conduct this second procedure with gender given the lack of relation between gender and either 1) pimfs incidence (in this sample; **Results**) or 2) reasoning, both in this sample (**Materials and Methods**) and in a larger sample (N = 523) (Wendelken et al. 2017).

T-tests were implemented with the t.test function from the R stats package. T-test effect sizes are reported with Cohen’s d (d) metric. The median and 95% confidence intervals were calculated with the MedianCI function from the DescTools R package. Linear regressions were run with the lm function from the stats package. Variable-ratio matching was implemented with the matchit function from the MatchIt package.

### Analysis IV: Control behavioral analyses

To ascertain whether the observed relationship between sulcal morphology and cognition is specific to reasoning performance, or generalizable to other measures of cognitive processing, we tested this sulcal-behavior relationship with measures of processing speed (Pattern Comparison Processing Speed Test) and list-sorting working memory (List Sorting Working Memory Test). Participant age and gender were not considered in these analyses, as they were not reliably associated with these tests of processing speed (*age*: ß = −0.05, *p* = 0.18; *gender*: χ^2^ = 0.05, *p* = 0.84) or working memory (*age*: ß = −0.001, *p* = 0.98; *gender*: χ^2^ = 1.58, *p* = 0.62). Two-sample t-tests were run to test for differences in performance on each measure as a function of left pimfs-v presence (*present* vs *absent*). We then used the Akaike Information Criterion (AIC) to compare the model predictions to reasoning predictions. Briefly, the AIC provides an estimate of in-sample prediction error and is suitable for non-nested model comparison. By comparing AIC scores, we are able to assess the relative performance of the two models. If the ΔAIC is >2, it suggests an interpretable difference between models. If the ΔAIC is >10, it suggests a strong difference between models, with the lower AIC value indicating the preferred model (Wagenmakers and Farrell 2004; Burnham and Anderson 2004).

Given that there was also a significant correlation between the Penn Matrices and List Sorting test scores (correlation matrix between all three behavioral variables in **Supplementary Fig. 4A**), and that List Sorting performance varied as a function of left pimfs-v sulcal presence/absence (**Supplementary Fig. 3A**), we tested whether this relationship was mediated by reasoning performance, consistent with the hypothesis that the behavioral-sulcal association for List Sorting is explained by relational reasoning demands of this specific working memory task. To this end, we implemented a bootstrapped causal mediation analysis (1,000 simulations) to quantify the average causal mediation effect (ACME) of reasoning on the left pimfs-v presence and WM relationship (mediation diagram in **Supplementary Fig. 4B**).

T-tests were implemented with the t.test function from the R stats package. T-test effect sizes are reported with Cohen’s d (d) metric. AIC values were quantified with the AIC function from the stats R package. The causal mediation analysis was implemented with the mediate function from the mediation package.

### Analysis V: Relating the size of the pimfs to reasoning performance

As in prior work (Clark et al. 2010; Garrison et al. 2015; Cachia et al. 2018; Rollins et al. 2020; Willbrand et al. 2022c), to determine whether it was truly a discrete sulcal metric that mattered, we followed up by testing whether a related continuous metric—the size, i.e., normalized surface area, of the pimfs—was also related to reasoning performance. To do so, we implemented a multiple linear regression with normalized total surface area of the pimfs component(s) in left and right hemispheres as predictors. Age and gender were not included, as they were not related to pimfs surface area (**Results**) or to reasoning performance. Linear regressions were implemented with the lm function from the stats package.

## Results

The pimfs was variably present within the 144 younger adult hemispheres examined (four example hemispheres depicting different types of pimfs patterning are presented in **Fig. 1A**). It was more common for participants to have two components in a given hemisphere (*left*: 72.22% of participants; *right*: 77.78%) than either one (*left*: 25%; *right*: 20.83%) or none (*left*: 2.78%; *right*: 1.39%; χ^2^s > 54, *p*s < 1.59e-12 in both hemispheres). When only one pimfs component was present, it was equally likely to be a dorsal or ventral component (χ^2^s < 2, *p*s > .31 in both hemispheres; **Fig. 1B**). The number of pimfs components and the presence of the pimfs-d and pimfs-v did not differ between hemispheres (*p*s > .23; **Fig. 1B**) or by participant age and gender (*p*s > .23); nor did the surface area of the pimfs differ by these features (*p*s > .60). There were no significant interactions between these variables in any analysis (*p*s > .23). These incidence rates were similar to those observed in our previous sample of children and adolescents (Willbrand et al. 2022c). To aid future identification in other samples, we share probabilistic maps of the pimfs components (**Fig. 1C**).

Since the pimfs was variably present among younger adults, we statistically tested whether this variability was related to reasoning performance, as previously found for children and adolescents using a similar matrix reasoning task (Willbrand et al. 2022c). The presence of two pimfs components in the left hemisphere was associated with 21% better reasoning performance on average (mean ± std = 18.1 ± 4.21) relative to either one or none (mean ± std = 15.0 ± 5.59; t(69) = 2.54, *p* = 0.026, *d* = 0.67). This effect was not driven by the fact that individuals with two components tended to have a larger overall pimfs surface area, since pimfs surface area (normalized by hemisphere surface area) was not related to reasoning (*p*sl J> 0.90; **Supplementary Fig. 1**). As previously found in children and adolescents (Willbrand et al. 2022c), this effect was driven by the presence or absence of the left pimfs-v: participants with a left pimfs-v (mean ± std = 18.1 ± 4.18) had on average 34% higher reasoning scores relative to those without it (mean ± std = 13.5 ± 5.57; t(69) = 3.44, *p* = 0.004, *d* = 1.03; **Fig. 2A**). Neither the right pimfs-v nor pimfs-d showed this effect (ts < 1.32, *p*s > 0.38, *d*s < 0.47). To account for the difference in sample sizes between adults with and without the left pimfs-v, we iteratively sampled a size-matched subset of the left pimfs-v present group 1,000 times. This procedure confirmed the behavioral difference (median [95% CI] *d* □=□0.92 [0.90-0.94]; **Fig. 2B**). Finally, although reasoning performance was unrelated to age in this younger adult sample (ß = −0.01, *p* = 0.98), we conducted additional analyses confirming that age did not confound the results (**Supplementary Results**; **Supplementary Fig. 2**).

Finally, to assess the generalizability and/or specificity of this brain-behavior relationship, we tested whether left pimfs-v presence was associated with performance on the Pattern Comparison Processing Speed Test or the List Sorting Working Memory Test, as processing speed and working memory are foundational cognitive skills that support reasoning (Kail and Salthouse 1994; Fry and Hale 2000; Ferrer et al. 2013; Kail et al. 2016; Holyoak and Monti 2021). The List Sorting test requires reordering items according to their relative size, thus requiring relational reasoning, in the form of transitive inference, in addition to working memory (**Materials and Methods**). Participants with a left pimfs-v (mean ± std = 114 ± 12.3) had on average 9% better performance on the List Sorting test on average relative to those without it (mean ± std = 105 ± 11.5; t(69) = 2.42, *p* = 0.035, *d* = 0.72; **Supplementary Fig. 3A**). While significant, this effect was not as large as for the Penn Matrices task task (ΔAIC_(working memory-reasoning)_ = 142.23). Additionally, the relationship between left pimfs-v presence and List Sorting scores was significantly mediated by Penn Matrices scores (via an indirect effect computed for 1,000 bootstrapped samples; average causal mediation effect [95% CI] = 6.13 [1.88, 10.88], *p* = 0.006; **Supplementary Fig. 4B**), further indicating that the relationship between left pimfs-v and relational reasoning is the stronger one. By contrast, left pimfs-v presence was not related to processing speed test performance (t(69) = −0.24, *p* = 0.81, *d* = −0.07; **Supplementary Fig. 3B**).

## Discussion

The pimfs exhibits prominent variability in humans that is robustly linked to variability in relational reasoning performance across age groups: in young adulthood, as reported here, and in childhood and adolescence (ages 6-18), as reported previously (Willbrand et al. 2022c). The pattern of results across the two studies was strikingly similar, with left pimfs-v–but not right pimfs-v or left or right pimfs-d—implicated in a test of matrix reasoning. We found that children and adults with left pimfs-v had on average 28% and 34% higher reasoning scores, respectively, than those without it. In the same sample of younger adults, we recently showed that pimfs-v and pimfs-d exhibited different patterns of large-scale resting-state functional connectivity, whereby the pimfs-v was associated with control-related networks and the pimfs-d was associated with attention-related networks (Willbrand et al. 2023a). Thus, the present behavioral dissociation between these sulcal components likely stems from participation in different brain networks and cognitive functions. In addition to showing that the sulcal-behavioral relationship did not generalize to all sulci, we showed that it did not generalize to all cognitive tasks. In both the adult and pediatric samples, the incidence of left pimfs-v was related to a test of relational reasoning but not processing speed—a finding suggesting some degree of specificity in this anatomical-behavior relationship. Notably, the studies covered different age ranges—ages 22-36 vs. 6-18—and also used different cognitive measures [our previous study used WISC-IV Matrix Reasoning task (Wechsler 1949) and the Woodcock-Johnson Psychoeducational Battery Cross Out task (Brown et al. 2012)]. Thus, taken together, these findings establish a robust anatomical-behavioral association that does not generalize to all forms of cognition or to all sulci, which converges with the observed functional dissociation between neighboring sulcal components (Willbrand et al. 2023a).

In addition to the abovementioned cognitive measures, both studies also included tests of working memory. Children and adolescents completed a standard task involving repeating a series of numbers in order or in reverse order [WISC-IV Digit Span WM task (Wechsler 1949)]. By contrast, adults completed a novel List Sorting task that requires participants to reorder a series of WM representations according to their size in real life. Size ordering, an example of transitive inference, is recognized as a form of relational reasoning (Halford et al. 1998). We found an effect of left pimfs-v presence on List Sorting in adults, but not on Digit Span in children. We posit, based in part on prior fMRI findings (Wendelken et al. 2008a), that the apparent discrepancy between these findings relates to substantive differences in task demands: namely, that List Sorting requires relational reasoning. This hypothesis remains to be tested.

The anatomical-behavioral link reported here and in our previous study (Willbrand et al. 2022c) appears to provide converging evidence with fMRI research on relational reasoning. Neurosynth, a meta-analytic tool drawing on over 14,000 fMRI studies (Yarkoni et al. 2011), shows that the most probable location of the pimfs-v includes, and forms the dorsal border of, a functionally defined area in aLPFC often referred to as rostrolateral prefrontal cortex that has been implicated in higher relational reasoning (Christoff et al. 2001; Kroger et al. 2002; Bunge et al. 2005; Wendelken et al. 2008a, b; Crone et al. 2009; Wendelken and Bunge 2010; Dumontheil et al. 2010; Krawczyk 2012; Vendetti and Bunge 2014; Hobeika et al. 2016; Hartogsveld et al. 2018; Assem et al. 2020; Holyoak and Monti 2021) (**Supplementary Fig. 5**). Future research should investigate this intriguing overlap, directly testing the spatial correspondence between the pimfs-v and reasoning-related task activations in individual participants. Future research should also address why, given these converging lines of evidence, some neuropsychological studies show that damage to aLPFC affects reasoning task accuracy (Urbanski et al. 2016; Bendetowicz et al. 2018) while others have not (Burgess 2000; Tranel et al. 2008; Waechter et al. 2013).

While we focus on the pimfs in the present study, and do show a sizeable relationship to high-level cognitive performance, we hypothesize that variable morphology of this sulcus reflects and/or drives neural differences more broadly. Indeed, the presence/absence and morphology of sulci are theorized to be anatomically linked to cortical white matter development (Sanides 1962, 1964; Van Essen et al. 2014; Reveley et al. 2015; Van Essen 2020; Cottaar et al. 2021; Miller et al. 2021b; Cachia et al. 2021), and the presence of sulci relates to changes in the local cytoarchitectonic organization of gray matter (Amiez et al. 2021). Neural efficiency is the foundation of our hypothesis regarding relationships between sulcal anatomy and reasoning observed in these and other studies (Voorhies et al. 2021; Willbrand et al. 2023b). Specifically, we (and others; Garrison et al., 2015) hypothesize that these sulcal-behavioral relationships stem from individual differences in white matter projections and, in turn, the distributed functional brain networks that underlie higher-level cognition. Future research should investigate this multiscale, mechanistic relationship describing the neural correlates of reasoning, integrating structural, functional, and behavioral data.

Given that the pimfs is rare in non-human hominoids (Hathaway et al. 2022; Amiez et al. 2023), the presence of left pimfs-v could reflect evolutionarily expanded white and gray matter properties that enhance neural communication in this higher cognitive area (Van Essen 1997, 2020; White et al. 2010; Zilles et al. 2013). Additionally, given (i) evidence implicating sulcal incidence to cognition in chimpanzees (Hopkins et al. 2021) and (ii) hypotheses that the disproportionate expansion of aLPFC in humans compared to non-human primates contributes to species differences in reasoning capacity (Semendeferi et al. 2001; Vendetti and Bunge 2014), another testable hypothesis is whether the incidence of the pimfs is also cognitively relevant in chimpanzees.

While the cognitive relevance of the pimfs has only been examined in neurotypical populations (Voorhies et al. 2021; Yao et al. 2022; Willbrand et al. 2022c), numerous studies have shown that variations in sulcal incidence are clinically relevant (Yücel et al. 2002, 2003; Le Provost et al. 2003; Fornito et al. 2006, 2008; Shim et al. 2009; Meredith et al. 2012; Gay et al. 2017; Nakamura et al. 2020; Harper et al. 2022). Thus, the present results raise the question: Does the incidence and/or morphology of the pimfs differ in clinical populations exhibiting impaired reasoning? Schizophrenia is a prime candidate for future investigations, given that it is marked by impaired reasoning (Weickert et al. 2000; Bowie and Harvey 2006; Keefe and Harvey 2012; Zhang et al. 2017; Alkan et al. 2021; McCutcheon et al. 2023) and has repeatedly been associated with altered aLPFC structure and function (Barnes et al. 2011; Tu et al. 2012; Kaplan et al. 2016; Kang et al. 2018; Pillinger et al. 2019; Nazli et al. 2020; Shinba et al. 2022). To help guide future studies examining the cognitive, evolutionary, developmental, clinical, and functional relevance of the pimfs, we share probabilistic predictions of the pimfs from our data (**Fig. 1C**).

In conclusion, we have shown that left pimfs-v presence is cognitively relevant in young adulthood, which extends previous work showing that left pimfs-v presence is cognitively relevant in childhood and adolescence. The combination of findings across studies establishes pimfs-v presence/absence as a novel developmental, cognitive, and evolutionarily relevant feature that should be considered in future studies in neurotypical and clinical populations examining how the complex relationships among multiscale anatomical and functional features of the brain give rise to abstract thought.

## Competing interests statement

The authors declare no competing financial interests.

## Data accessibility statement

Data, code, analysis pipelines, and sulcal probability maps, are on GitHub (https://github.com/cnl-berkeley/stable_projects/tree/main/CognitiveRelevance_PrefrontalStructure).

## Author contributions statement

EHW, SAB, and KSW designed research; EHW, SJ, SC, CBH, WIV, and KSW performed manual sulcal labeling; EHW, SAB, and KSW analyzed data; EHW, SAB, and KSW wrote the paper; all authors gave final approval to the paper before submission.

## Acknowledgments

This research was supported by NICHD R21HD100858 (Weiner, Bunge), an NSF CAREER Award 2042251 (Weiner), and an NSF-GRFP fellowship (Voorhies). Young adult neuroimaging and behavioral data were provided by the HCP, WU-Minn Consortium (Principal Investigators: David Van Essen and Kamil Ugurbil; NIH Grant 1U54-MH-091657) funded by the 16 NIH Institutes and Centers that support the NIH Blueprint for Neuroscience Research, and the McDonnell Center for Systems Neuroscience at Washington University. We thank Jewelia Yao and Jacob Miller for their assistance defining other additional lateral prefrontal cortex sulci across age groups. We also thank the HCP researchers for participant recruitment and data collection and sharing, as well as the participants who took part in the study.

## Notes

### Competing Interest Statement

The authors have declared no competing interest.

### Summary of Updates

Updates to main text and supplements; changed title

## References

Alexander PA (2016) Relational thinking and relational reasoning: harnessing the power of patterning. NPJ Sci Learn 1:16004. 10.1038/npjscilearn.2016.4

Alkan E, Davies G, Evans SL (2021) Cognitive impairment in schizophrenia: relationships with cortical thickness in fronto-temporal regions, and dissociability from symptom severity. NPJ Schizophr 7:20. 10.1038/s41537-021-00149-0

Allen M, Poggiali D, Whitaker K, et al (2021) Raincloud plots: a multi-platform tool for robust data visualization. Wellcome Open Res 4:63. 10.12688/wellcomeopenres.15191.2

Amiez C, Sallet J, Giacometti C, et al (2023) A revised perspective on the evolution of the lateral frontal cortex in primates. Science Advances 9:eadf9445. 10.1126/sciadv.adf9445

Amiez C, Sallet J, Hopkins WD, et al (2019) Sulcal organization in the medial frontal cortex provides insights into primate brain evolution. Nat Commun 10:1–14. 10.1038/s41467-019-11347-x

Amiez C, Sallet J, Novek J, et al (2021) Chimpanzee histology and functional brain imaging show that the paracingulate sulcus is not human-specific. Commun Biol 4:54. 10.1038/s42003-020-01571-3

Assem M, Glasser MF, Van Essen DC, Duncan J (2020) A Domain-General Cognitive Core Defined in Multimodally Parcellated Human Cortex. Cereb Cortex 30:4361–4380. 10.1093/cercor/bhaa023

Barch DM, Burgess GC, Harms MP, et al (2013) Function in the human connectome: task-fMRI and individual differences in behavior. Neuroimage 80:169–189. 10.1016/j.neuroimage.2013.05.033

Barnes MR, Huxley-Jones J, Maycox PR, et al (2011) Transcription and pathway analysis of the superior temporal cortex and anterior prefrontal cortex in schizophrenia. J Neurosci Res 89:1218–1227. 10.1002/jnr.22647

Bendetowicz D, Urbanski M, Garcin B, et al (2018) Two critical brain networks for generation and combination of remote associations. Brain 141:217–233. 10.1093/brain/awx294

Benson NC, Butt OH, Datta R, et al (2012) The retinotopic organization of striate cortex is well predicted by surface topology. Curr Biol 22:2081–2085. 10.1016/j.cub.2012.09.014

Boling W, Olivier A, Bittar RG, Reutens D (1999) Localization of hand motor activation in Broca’s pli de passage moyen. J Neurosurg 91:903–910. 10.3171/jns.1999.91.6.0903

Bowie CR, Harvey PD (2006) Cognitive deficits and functional outcome in schizophrenia. Neuropsychiatr Dis Treat 2:531–536. 10.2147/nedt.2006.2.4.531

Brown TT, Kuperman JM, Chung Y, et al (2012) Neuroanatomical assessment of biological maturity. Curr Biol 22:1693–1698. 10.1016/j.cub.2012.07.002

Bunge SA, Wendelken C, Badre D, Wagner AD (2005) Analogical reasoning and prefrontal cortex: evidence for separable retrieval and integration mechanisms. Cereb Cortex 15:239–249. 10.1093/cercor/bhh126

Burgess PW (2000) Strategy application disorder: the role of the frontal lobes in human multitasking. Psychol Res 63:279–288. 10.1007/s004269900006

Burnham KP, Anderson DR (2004) Multimodel Inference: Understanding AIC and BIC in Model Selection. Sociol Methods Res 33:261–304. 10.1177/0049124104268644

Cachia A, Borst G, Jardri R, et al (2021) Towards Deciphering the Fetal Foundation of Normal Cognition and Cognitive Symptoms From Sulcation of the Cortex. Front Neuroanat 15:712862. 10.3389/fnana.2021.712862

Cachia A, Roell M, Mangin J-F, et al (2018) How interindividual differences in brain anatomy shape reading accuracy. Brain Struct Funct 223:701–712. 10.1007/s00429-017-1516-x

Carpenter PA, Just MA, Shell P (1990) What one intelligence test measures: a theoretical account of the processing in the Raven Progressive Matrices Test. Psychol Rev 97:404–431. 10.1037/0033-295X.97.3.404

Cattell RB (1943) The measurement of adult intelligence. Psychol Bull 40:153–193. 10.1037/h0059973

Christoff K, Prabhakaran V, Dorfman J, et al (2001) Rostrolateral prefrontal cortex involvement in relational integration during reasoning. Neuroimage 14:1136–1149. 10.1006/nimg.2001.0922

Clark GM, Mackay CE, Davidson ME, et al (2010) Paracingulate sulcus asymmetry; sex difference, correlation with semantic fluency and change over time in adolescent onset psychosis. Psychiatry Res 184:10–15. 10.1016/j.pscychresns.2010.06.012

Cottaar M, Bastiani M, Boddu N, et al (2021) Modelling white matter in gyral blades as a continuous vector field. Neuroimage 227:117693. 10.1016/j.neuroimage.2020.117693

Crone EA, Wendelken C, van Leijenhorst L, et al (2009) Neurocognitive development of relational reasoning. Dev Sci 12:55–66. 10.1111/j.1467-7687.2008.00743.x

Cykowski MD, Coulon O, Kochunov PV, et al (2008) The central sulcus: an observer-independent characterization of sulcal landmarks and depth asymmetry. Cereb Cortex 18:1999–2009. 10.1093/cercor/bhm224

Dale AM, Fischl B, Sereno MI (1999) Cortical surface-based analysis. I. Segmentation and surface reconstruction. Neuroimage 9:179–194. 10.1006/nimg.1998.0395

Dumontheil I, Houlton R, Christoff K, Blakemore S-J (2010) Development of relational reasoning during adolescence. Dev Sci 13:F15–24. 10.1111/j.1467-7687.2010.01014.x

Ferrer E, Whitaker KJ, Steele JS, et al (2013) White matter maturation supports the development of reasoning ability through its influence on processing speed. Dev Sci 16:941–951. 10.1111/desc.12088

Fischl B, Dale AM (2000) Measuring the thickness of the human cerebral cortex from magnetic resonance images. Proc Natl Acad Sci U S A 97:11050–11055. 10.1073/pnas.200033797

Fischl B, Sereno MI, Dale AM (1999) Cortical surface-based analysis. II: Inflation, flattening, and a surface-based coordinate system. Neuroimage 9:195–207. 10.1006/nimg.1998.0396

Fornito A, Malhi GS, Lagopoulos J, et al (2008) Anatomical abnormalities of the anterior cingulate and paracingulate cortex in patients with bipolar I disorder. Psychiatry Research: Neuroimaging 162:123–132

Fornito A, Yücel M, Wood SJ, et al (2006) Morphology of the paracingulate sulcus and executive cognition in schizophrenia. Schizophr Res 88:192–197. 10.1016/j.schres.2006.06.034

Fry AF, Hale S (2000) Relationships among processing speed, working memory, and fluid intelligence in children. Biol Psychol 54:1–34. 10.1016/s0301-0511(00)00051-x

Garrison JR, Fernyhough C, McCarthy-Jones S, et al (2015) Paracingulate sulcus morphology is associated with hallucinations in the human brain. Nat Commun 6:8956. 10.1038/ncomms9956

Gay O, Plaze M, Oppenheim C, et al (2017) Cognitive control deficit in patients with first-episode schizophrenia is associated with complex deviations of early brain development. J Psychiatry Neurosci 42:87–94

Glasser MF, Sotiropoulos SN, Wilson JA, et al (2013) The minimal preprocessing pipelines for the Human Connectome Project. Neuroimage 80:105–124. 10.1016/j.neuroimage.2013.04.127

Gratton C, Nelson SM, Gordon EM (2022) Brain-behavior correlations: Two paths toward reliability. Neuron 110:1446–1449

Halford GS, Wilson WH, Phillips S (1998) Processing capacity defined by relational complexity: implications for comparative, developmental, and cognitive psychology. Behav Brain Sci 21:803–31; discussion 831–64. 10.1017/s0140525x98001769

Harper L, Lindberg O, Bocchetta M, et al (2022) Prenatal Gyrification Pattern Affects Age at Onset in Frontotemporal Dementia. Cereb Cortex. 10.1093/cercor/bhab457

Hartogsveld B, Bramson B, Vijayakumar S, et al (2018) Lateral frontal pole and relational processing: Activation patterns and connectivity profile. Behav Brain Res 355:2–11. 10.1016/j.bbr.2017.08.003

Hathaway CB, Voorhies WI, Sathishkumar N, et al (2023) Defining putative tertiary sulci in lateral prefrontal cortex in chimpanzees using human predictions. Brain Struct Funct. 10.1007/s00429-023-02638-7

Hathaway CB, Voorhies WI, Sathishkumar N, et al (2022) Defining tertiary sulci in lateral prefrontal cortex in chimpanzees using human predictions. bioRxiv 2022.04.12.488091

Hinds OP, Rajendran N, Polimeni JR, et al (2008) Accurate prediction of V1 location from cortical folds in a surface coordinate system. Neuroimage 39:1585–1599. 10.1016/j.neuroimage.2007.10.033

Hobeika L, Diard-Detoeuf C, Garcin B, et al (2016) General and specialized brain correlates for analogical reasoning: A meta-analysis of functional imaging studies. Hum Brain Mapp 37:1953–1969. 10.1002/hbm.23149

Holyoak KJ, Monti MM (2021) Relational Integration in the Human Brain: A Review and Synthesis. Journal of Cognitive Neuroscience 33:341–356

Hopkins WD, Procyk E, Petrides M, et al (2021) Sulcal Morphology in Cingulate Cortex is Associated with Voluntary Oro-Facial Motor Control and Gestural Communication in Chimpanzees (Pan troglodytes). Cereb Cortex 31:2845–2854. 10.1093/cercor/bhaa392

James W (1890a) The principles of psychology, Vol I

James W (1890b) The Principles Of Psychology Volume II By William James (1890)

Kail R, Salthouse TA (1994) Processing speed as a mental capacity. Acta Psychol 86:199–225. 10.1016/0001-6918(94)90003-5

Kail RV, Lervåg A, Hulme C (2016) Longitudinal evidence linking processing speed to the development of reasoning. Dev Sci 19:1067–1074. 10.1111/desc.12352

Kang SS, MacDonald AW 3rd, Chafee MV, et al (2018) Abnormal cortical neural synchrony during working memory in schizophrenia. Clin Neurophysiol 129:210–221. 10.1016/j.clinph.2017.10.024

Kaplan CM, Saha D, Molina JL, et al (2016) Estimating changing contexts in schizophrenia. Brain 139:2082–2095. 10.1093/brain/aww095

Keefe RSE, Harvey PD (2012) Cognitive Impairment in Schizophrenia. In: Geyer MA, Gross G (eds) Novel Antischizophrenia Treatments. Springer Berlin Heidelberg, Berlin, Heidelberg, pp 11–37

Krawczyk DC (2012) The cognition and neuroscience of relational reasoning. Brain Res 1428:13–23. 10.1016/j.brainres.2010.11.080

Kroger JK, Sabb FW, Fales CL, et al (2002) Recruitment of anterior dorsolateral prefrontal cortex in human reasoning: a parametric study of relational complexity. Cereb Cortex 12:477–485. 10.1093/cercor/12.5.477

Le Provost J-B, Bartres-Faz D, Paillere-Martinot M-L, et al (2003) Paracingulate sulcus morphology in men with early-onset schizophrenia. Br J Psychiatry 182:228–232. 10.1192/bjp.182.3.228

Li S, Han Y, Wang D, et al (2010) Mapping surface variability of the central sulcus in musicians. Cereb Cortex 20:25–33. 10.1093/cercor/bhp074

Marek S, Tervo-Clemmens B, Calabro FJ, et al (2022) Reproducible brain-wide association studies require thousands of individuals. Nature 603:654–660. 10.1038/s41586-022-04492-9

McArdle JJ, Ferrer-Caja E, Hamagami F, Woodcock RW (2002) Comparative longitudinal structural analyses of the growth and decline of multiple intellectual abilities over the life span. Dev Psychol 38:115–142. 10.1037/0012-1649.38.1.115

McCutcheon RA, Keefe RSE, McGuire PK (2023) Cognitive impairment in schizophrenia: aetiology, pathophysiology, and treatment. Mol Psychiatry. 10.1038/s41380-023-01949-9

Meredith SM, Whyler NCA, Stanfield AC, et al (2012) Anterior cingulate morphology in people at genetic high-risk of schizophrenia. Eur Psychiatry 27:377–385. 10.1016/j.eurpsy.2011.11.004

Miller JA, D’Esposito M, Weiner KS (2021a) Using Tertiary Sulci to Map the “Cognitive Globe” of Prefrontal Cortex. J Cogn Neurosci 1–18. 10.1162/jocn_a_01696

Miller JA, Voorhies WI, Li X, et al (2020) Sulcal morphology of ventral temporal cortex is shared between humans and other hominoids. Sci Rep 10:17132. 10.1038/s41598-020-73213-x

Miller JA, Voorhies WI, Lurie DJ, et al (2021b) Overlooked Tertiary Sulci Serve as a Meso-Scale Link between Microstructural and Functional Properties of Human Lateral Prefrontal Cortex. J Neurosci 41:2229–2244. 10.1523/JNEUROSCI.2362-20.2021

Ming K, Rosenbaum PR (2000) Substantial gains in bias reduction from matching with a variable number of controls. Biometrics 56:118–124. 10.1111/j.0006-341x.2000.00118.x

Nakamura M, Nestor PG, Shenton ME (2020) Orbitofrontal Sulcogyral Pattern as a Transdiagnostic Trait Marker of Early Neurodevelopment in the Social Brain. Clin EEG Neurosci 51:275–284. 10.1177/1550059420904180

Nazli ŞB, Koçak OM, Kirkici B, et al (2020) Investigation of the Processing of Noun and Verb Words with fMRI in Patients with Schizophrenia. Noro Psikiyatr Ars 57:9–14. 10.29399/npa.23521

Paus T, Tomaiuolo F, Otaky N, et al (1996) Human Cingulate and Paracingulate Sulci: Pattern, Variability, Asymmetry, and Probabilistic Map. Cerebral Cortex 6:207–214

Petrides M (2019) Atlas of the Morphology of the Human Cerebral Cortex on the Average MNI Brain. Academic Press

Petrides M (2013) Neuroanatomy of Language Regions of the Human Brain. Academic Press

Pillinger T, Rogdaki M, McCutcheon RA, et al (2019) Altered glutamatergic response and functional connectivity in treatment resistant schizophrenia: the effect of riluzole and therapeutic implications. Psychopharmacology 236:1985–1997. 10.1007/s00213-019-5188-5

Raven JC (1941) Standardization of progressive matrices, 1938. Br J Med Psychol 19:137–150. 10.1111/j.2044-8341.1941.tb00316.x

Reveley C, Seth AK, Pierpaoli C, et al (2015) Superficial white matter fiber systems impede detection of long-range cortical connections in diffusion MR tractography. Proc Natl Acad Sci U S A 112:E2820–8. 10.1073/pnas.1418198112

Rollins CPE, Garrison JR, Arribas M, et al (2020) Evidence in cortical folding patterns for prenatal predispositions to hallucinations in schizophrenia. Transl Psychiatry 10:387. 10.1038/s41398-020-01075-y

Sanides F (1962) Besprechung. In: Sanides F (ed) Die Architektonik des Menschlichen Stirnhirns: Zugleich eine Darstellung der Prinzipien Seiner Gestaltung als Spiegel der Stammesgeschichtlichen Differenzierung der Grosshirnrinde. Springer Berlin Heidelberg, Berlin, Heidelberg, pp 176–190

Sanides F (1964) Structure and function of the human frontal lobe. Neuropsychologia 2:209–219. 10.1016/0028-3932(64)90005-3

Semendeferi K, Armstrong E, Schleicher A, et al (2001) Prefrontal cortex in humans and apes: a comparative study of area 10. Am J Phys Anthropol 114:224–241. 10.1002/1096-8644(200103)114:3<224::AID-AJPA1022>3.0.CO;2-I

Shim G, Jung WH, Choi J-S, et al (2009) Reduced cortical folding of the anterior cingulate cortex in obsessive-compulsive disorder. J Psychiatry Neurosci 34:443–449

Shinba T, Kariya N, Matsuda S, et al (2022) Near-Infrared Time-Resolved Spectroscopy Shows Anterior Prefrontal Blood Volume Reduction in Schizophrenia but Not in Major Depressive Disorder. Sensors 22.: 10.3390/s22041594

Sun ZY, Klöppel S, Rivière D, et al (2012) The effect of handedness on the shape of the central sulcus. Neuroimage 60:332–339. 10.1016/j.neuroimage.2011.12.050

Tranel D, Manzel K, Anderson SW (2008) Is the prefrontal cortex important for fluid intelligence? A neuropsychological study using Matrix Reasoning. Clin Neuropsychol 22:242–261. 10.1080/13854040701218410

Tu P-C, Hsieh J-C, Li C-T, et al (2012) Cortico-striatal disconnection within the cingulo-opercular network in schizophrenia revealed by intrinsic functional connectivity analysis: a resting fMRI study. Neuroimage 59:238–247. 10.1016/j.neuroimage.2011.07.086

Urbanski M, Bréchemier M-L, Garcin B, et al (2016) Reasoning by analogy requires the left frontal pole: lesion-deficit mapping and clinical implications. Brain 139:1783–1799. 10.1093/brain/aww072

Van Essen DC (2020) A 2020 view of tension-based cortical morphogenesis. Proc Natl Acad Sci U S A. 10.1073/pnas.2016830117

Van Essen DC (2007) 4.16 - Cerebral Cortical Folding Patterns in Primates: Why They Vary and What They Signify. In: Kaas JH (ed) Evolution of Nervous Systems. Academic Press, Oxford, pp 267–276

Van Essen DC (1997) A tension-based theory of morphogenesis and compact wiring in the central nervous system. Nature 385:313–318. 10.1038/385313a0

Van Essen DC, Jbabdi S, Sotiropoulos SN, et al (2014) Mapping connections in humans and non-human primates. In: Diffusion MRI. Elsevier, pp 337–358

Vendetti MS, Bunge SA (2014) Evolutionary and developmental changes in the lateral frontoparietal network: a little goes a long way for higher-level cognition. Neuron 84:906– 917. 10.1016/j.neuron.2014.09.035

Voorhies WI, Miller JA, Yao JK, et al (2021) Cognitive insights from tertiary sulci in prefrontal cortex. Nat Commun 12:5122. 10.1038/s41467-021-25162-w

Waechter RL, Goel V, Raymont V, et al (2013) Transitive inference reasoning is impaired by focal lesions in parietal cortex rather than rostrolateral prefrontal cortex. Neuropsychologia 51:464–471. 10.1016/j.neuropsychologia.2012.11.026

Wagenmakers E-J, Farrell S (2004) AIC model selection using Akaike weights. Psychon Bull Rev 11:192–196. 10.3758/bf03206482

Wandell BA, Winawer J (2011) Imaging retinotopic maps in the human brain. Vision Res 51:718–737. 10.1016/j.visres.2010.08.004

Wechsler D (1949) Wechsler Intelligence Scale for Children; manual. 113:

Weickert TW, Goldberg TE, Gold JM, et al (2000) Cognitive impairments in patients with schizophrenia displaying preserved and compromised intellect. Arch Gen Psychiatry 57:907–913. 10.1001/archpsyc.57.9.907

Wei X, Yin Y, Rong M, et al (2017) Paracingulate Sulcus Asymmetry in the Human Brain: Effects of Sex, Handedness, and Race. Sci Rep 7:42033. 10.1038/srep42033

Wendelken C, Bunge SA (2010) Transitive inference: distinct contributions of rostrolateral prefrontal cortex and the hippocampus. J Cogn Neurosci 22:837–847. 10.1162/jocn.2009.21226

Wendelken C, Bunge SA, Carter CS (2008a) Maintaining structured information: an investigation into functions of parietal and lateral prefrontal cortices. Neuropsychologia 46:665–678. 10.1016/j.neuropsychologia.2007.09.015

Wendelken C, Ferrer E, Ghetti S, et al (2017) Frontoparietal Structural Connectivity in Childhood Predicts Development of Functional Connectivity and Reasoning Ability: A Large-Scale Longitudinal Investigation. J Neurosci 37:8549–8558. 10.1523/JNEUROSCI.3726-16.2017

Wendelken C, Nakhabenko D, Donohue SE, et al (2008b) “Brain is to thought as stomach is to ??”: investigating the role of rostrolateral prefrontal cortex in relational reasoning. J Cogn Neurosci 20:682–693. 10.1162/jocn.2008.20055

Westlin C, Theriault JE, Katsumi Y, et al (2023) Improving the study of brain-behavior relationships by revisiting basic assumptions. Trends Cogn Sci. 10.1016/j.tics.2022.12.015

White T, Su S, Schmidt M, et al (2010) The development of gyrification in childhood and adolescence. Brain Cogn 72:36–45. 10.1016/j.bandc.2009.10.009

Willbrand EH, Bunge SA, Weiner KS (2023a) Neuroanatomical and functional dissociations between variably present anterior lateral prefrontal sulci. bioRxiv 2023.05.25.542301

Willbrand EH, Ferrer E, Bunge SA, Weiner KS (2023b) Development of human lateral prefrontal sulcal morphology and its relation to reasoning performance. J Neurosci. 10.1523/JNEUROSCI.1745-22.2023

Willbrand EH, Ferrer E, Bunge SA, Weiner KS (2022a) Development of human lateral prefrontal sulcal morphology and its relation to reasoning performance. bioRxiv 2022.09.14.507822

Willbrand EH, Maboudian SA, Kelly JP, et al (2023c) Sulcal morphology of posteromedial cortex substantially differs between humans and chimpanzees. Communications Biology 6:1–14. 10.1038/s42003-023-04953-5

Willbrand EH, Parker BJ, Voorhies WI, et al (2022b) Uncovering a tripartite landmark in posterior cingulate cortex. Science Advances 8:eabn9516. 10.1126/sciadv.abn9516

Willbrand EH, Voorhies WI, Yao JK, et al (2022c) Presence or absence of a prefrontal sulcus is linked to reasoning performance during child development. Brain Struct Funct 227:2543–2551. 10.1007/s00429-022-02539-1

Yao JK, Voorhies WI, Miller JA, et al (2022) Sulcal depth in prefrontal cortex: a novel predictor of working memory performance. Cereb Cortex bhac173. 10.1093/cercor/bhac173

Yarkoni T, Poldrack RA, Nichols TE, et al (2011) Large-scale automated synthesis of human functional neuroimaging data. Nat Methods 8:665–670. 10.1038/nmeth.1635

Yousry TA, Schmid UD, Alkadhi H, et al (1997) Localization of the motor hand area to a knob on the precentral gyrus. A new landmark. Brain 120 (Pt 1):141–157. 10.1093/brain/120.1.141

Yücel M, Stuart GW, Maruff P, et al (2002) Paracingulate morphologic differences in males with established schizophrenia: a magnetic resonance imaging morphometric study. Biol Psychiatry 52:15–23. 10.1016/s0006-3223(02)01312-4

Yücel M, Wood SJ, Phillips LJ, et al (2003) Morphology of the anterior cingulate cortex in young men at ultra-high risk of developing a psychotic illness. Br J Psychiatry 182:518–524. 10.1192/bjp.182.6.518

Zhang B, Han M, Tan S, et al (2017) Gender differences measured by the MATRICS consensus cognitive battery in chronic schizophrenia patients. Sci Rep 7:11821. 10.1038/s41598-017-12027-w

Zilles K, Armstrong E, Schleicher A, Kretschmann H-J (1988) The human pattern of gyrification in the cerebral cortex. Anatomy and Embryology 179:173–179

Zilles K, Palomero-Gallagher N, Amunts K (2013) Development of cortical folding during evolution and ontogeny. Trends Neurosci 36:275–284. 10.1016/j.tins.2013.01.006

