## Supplementary Material for "Sulcal variability in anterior lateral prefrontal cortex contributes to variability in reasoning performance among young adults"

Cognitive relevance of an evolutionarily new and variable prefrontal structure

**Authors and affiliations**

**Co-senior authors*

**Supplementary Results**

**Relationship between reasoning performance and left pimfs-v presence is not confounded by age**

To confirm that the relationship between left pimfs-v presence and reasoning performance was not confounded by age, we implemented a 3-step procedure (**Materials and Methods**). First, a linear regression with left pimfs-v presence and age as predictors in the full sample showed that left pimfs-v presence was still a significant predictor of reasoning performance when controlling for age (ß = 4.57, *t* = 3.40, *p* = 0.001). Further, this model explained significantly more variance in reasoning (adjusted R^2^ = 0.12, *p* = .004) compared to a model with age alone (adjusted R^2^  < .01, *p* = .74; model comparison: *p* = 0.001).

We then further confirmed the aforementioned results by employing variable-ratio matching (**Materials and Methods**) to create an age-matched sample with the original 14 participants without a left pimfs-v and 42 age-matched participants with a left pimfs-v (mean age = 29.92, eCDF). A weighted regression with this matched sample (with left pimfs-v and age as predictors) revealed that left pimfs-v presence remained a significant predictor (ß = 4.81, *t* = 3.29, *p* = 0.001; **Supplementary Fig. 2**). Critically, this model also explained significantly more variance in reasoning (adjusted R^2^ = 0.13, *p* = .007) compared to a model with age alone (adjusted R^2^  < .01, *p* = .89; model comparison: *p* = 0.001). We further evaluated the model fit with repeated K-fold cross-validation (five-fold, 10 repeats). The model including left pimfs-v presence as a factor showed increased prediction accuracy and decreased RMSE_CV_ (R^2^_CV_ = 0.20, RMSE_CV_ = 4.66) compared to a model with age only (R^2^_CV_ = 0.12, RMSE_CV_ = 5.04). Altogether, these results indicate that the presence of the left pimfs-v explained unique variance in reasoning scores above and beyond age.

**Supplementary Figures**

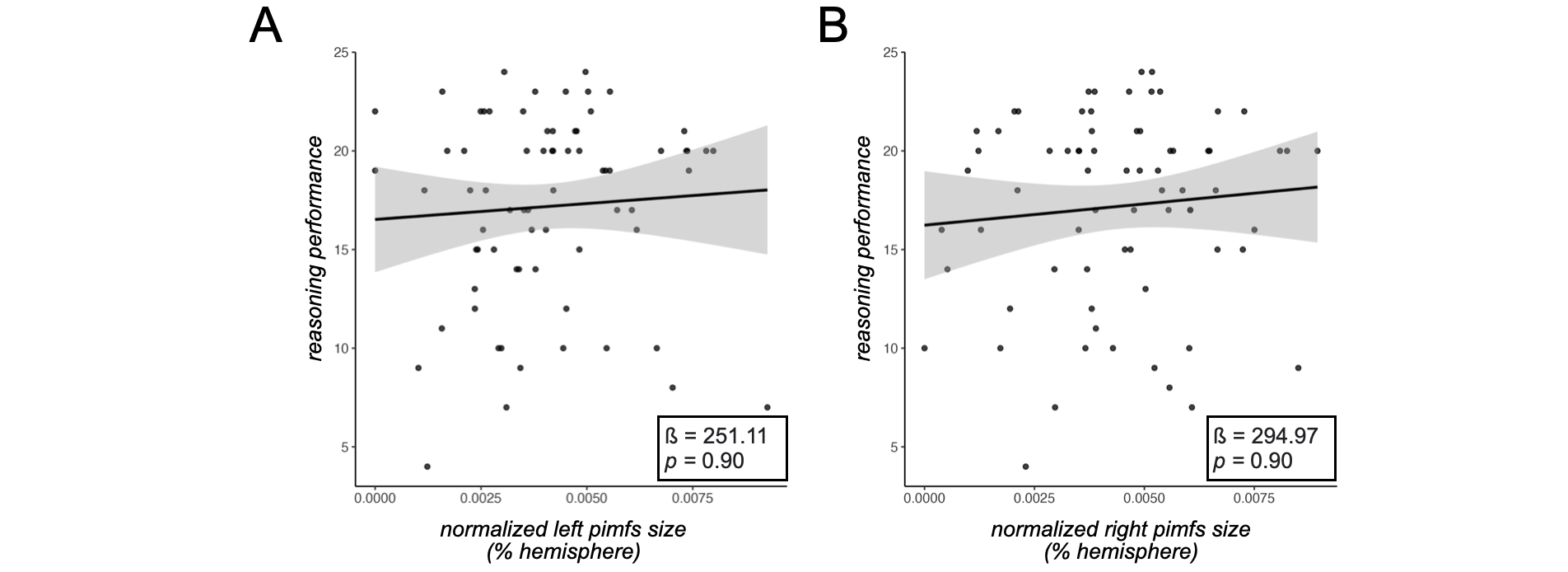

**Supplementary Figure 1. Reasoning performance is unrelated to pimfs size. A.** Scatterplot of reasoning performance as a function of left pimfs size (quantified as left pimfs surface area in mm^2^, normalized by left hemisphere surface area in mm^2^). Dots indicate values for each individual participant. The best fit line ± 95% confidence interval is also included along with the beta-value (ß) and *p*-value (FDR corrected) from the linear regression. **B.** Same format as A for right pimfs size.

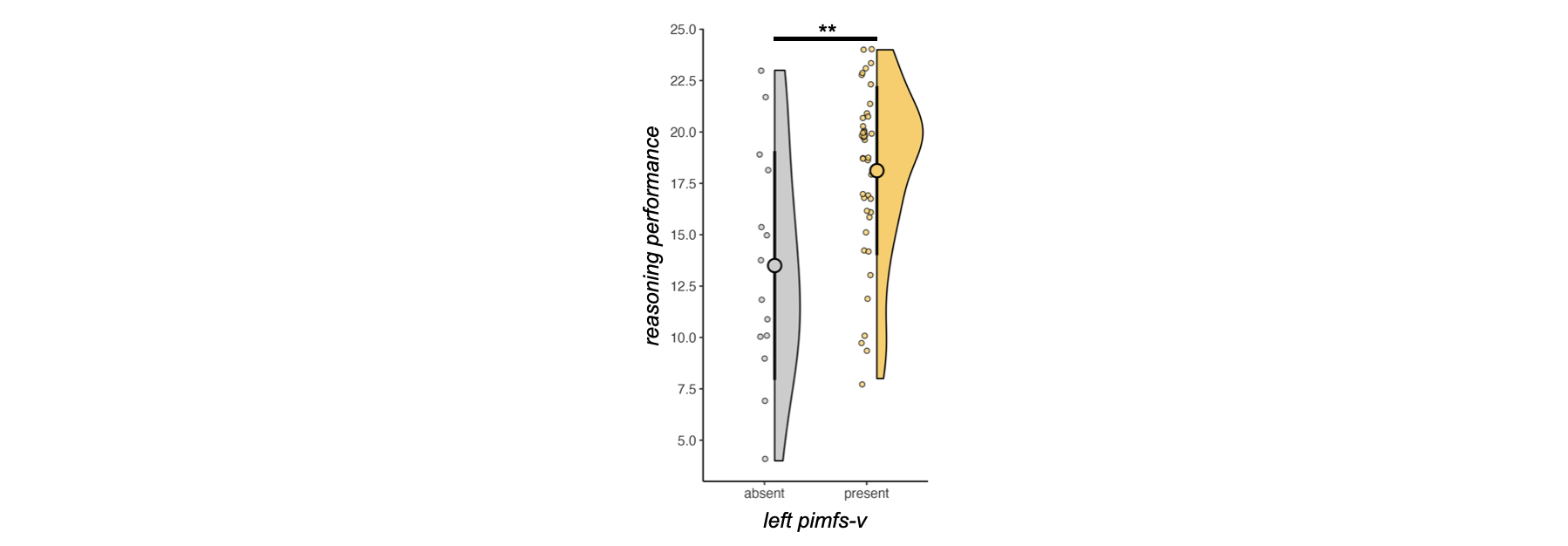

**Supplementary figure 2. Reasoning performance is related to left pimfs-v presence in the matched sample.** Raincloud plots depicting Penn Progressive Matrices task score as a function of left pimfs-v presence in younger adults (present, N = 42; absent, N = 14). Large dots and error bars represent mean ± std reasoning score; violin plots represent kernel density estimate. Small dots indicate individual participants. (** *p* < 0.01).

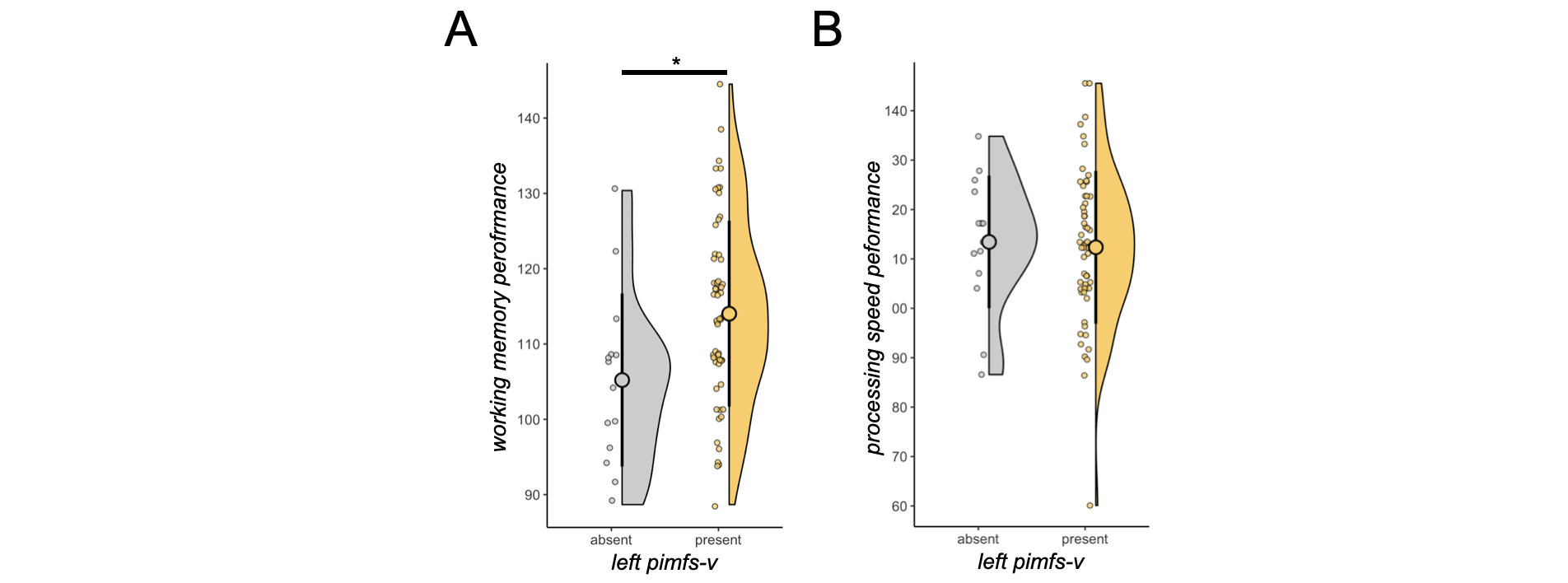
**Supplementary Figure 3. List-sorting working memory performance, but not processing speed performance, is related to left pimfs-v presence. A.** Same format as supplementary figure 2 for the List Sorting task in the full sample (N = 71). **B.** Same format as A for the Pattern Completion task. (* *p* < 0.05; no asterisks *p* > 0.05).

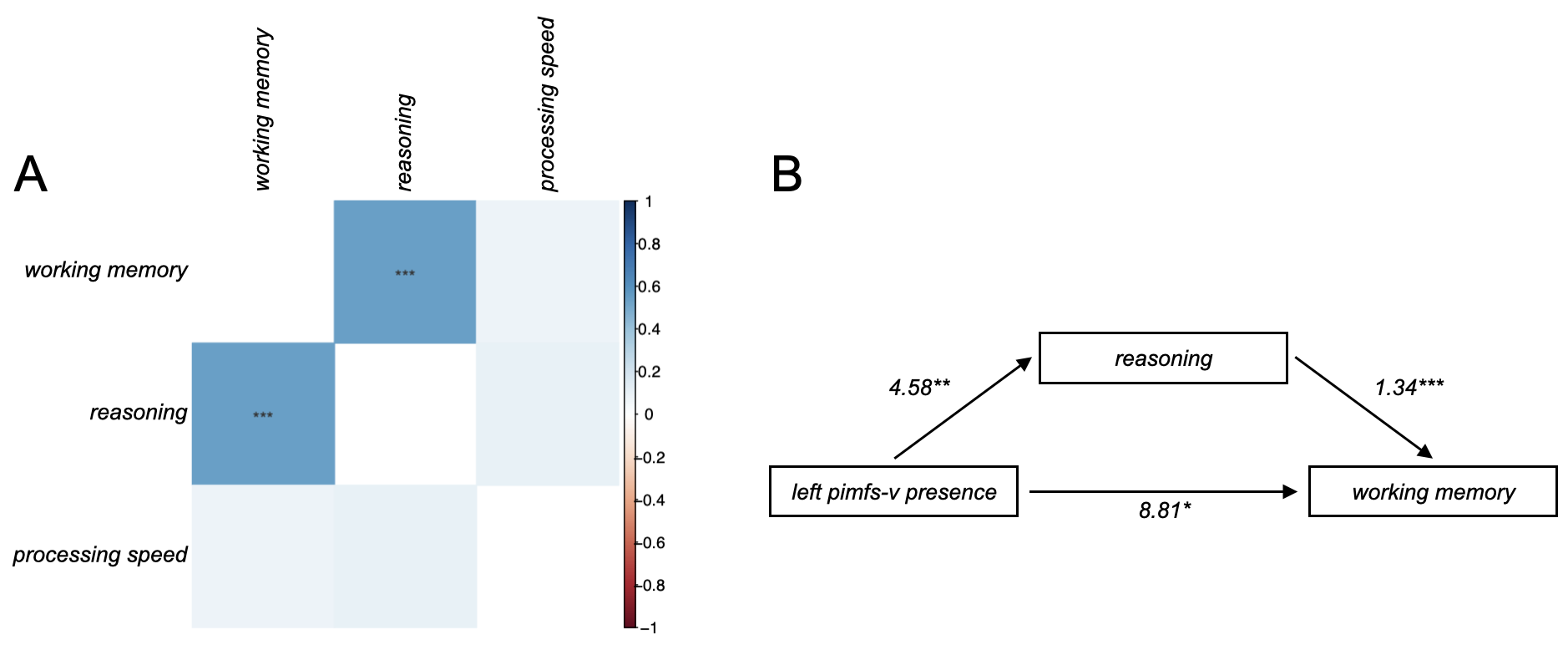
**Supplementary Figure 4. Relationship between list-sorting working memory performance and left pimfs-v presence is mediated by reasoning performance. A.** Three-by-three correlation matrix displaying the correlation coefficient (colors; see scale) and *p*-values for each correlation between the three behavioral variables. **B.** Mediation diagram showing that the relationship between left pimfs-v presence and working memory is significantly mediated by reasoning performance (via an indirect effect computed for 1,000 bootstrapped samples; average causal mediation effect [95% CI] = 6.13 [1.88, 10.88], *p* = 0.006). Beta-values (numbers) and *p*-values (asterisks) are included for each relationship. (*** *p* < 0.001; ** *p* < 0.01; * *p* < 0.05; no asterisks *p* > 0.05).

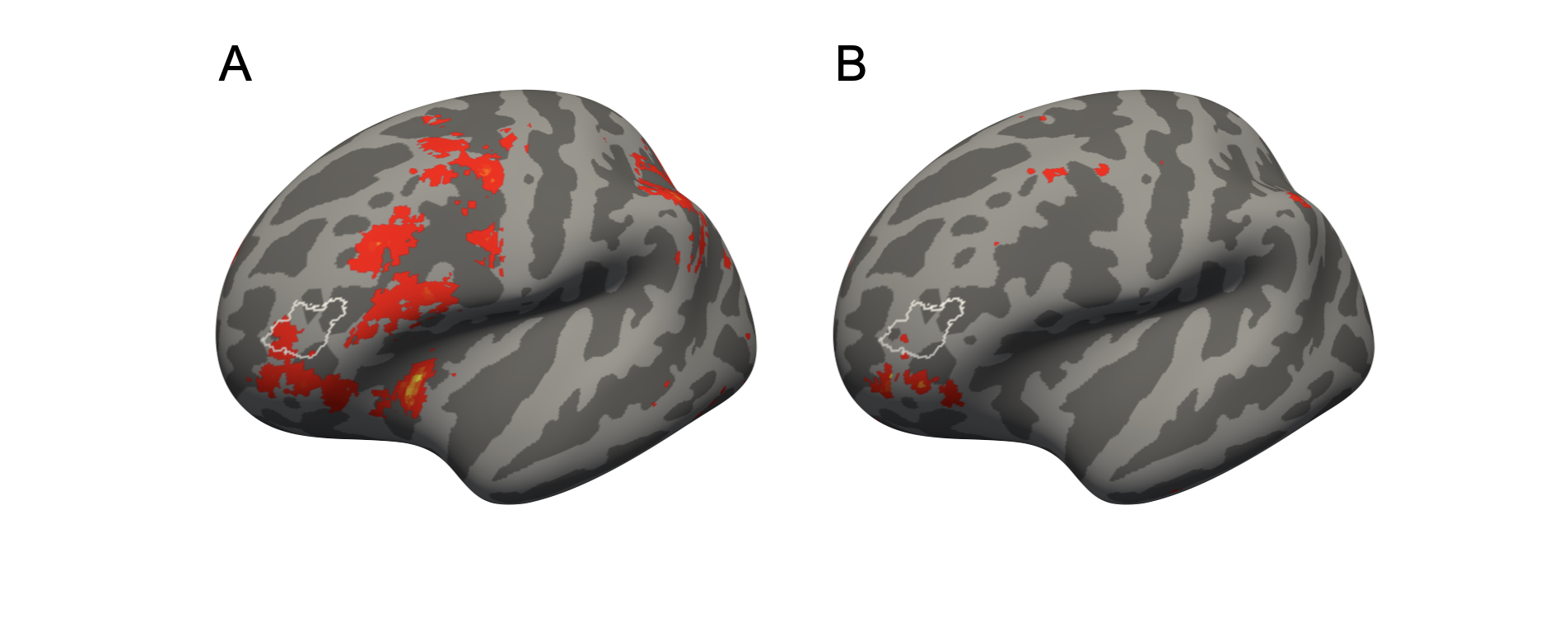
**Supplementary Figure 5. A Neurosynth meta-analysis shows that the most probable location of the left pimfs-v includes and forms the dorsal border of a functional region that has been preferentially implicated in reasoning. A.** Left hemisphere inflated *fsaverage* cortical surface displaying overlap of a whole-brain FDR-corrected (*p* = 0.01) uniformity-test meta-analysis z-score map of the “reasoning” term (red heatmap; downloaded from Neurosynth (Yarkoni et al. 2011) (https:// neurosynth.org/) and the left pimfs-v MPM (white outline; from Fig. 1C). This map was generated from a χ^2^ test comparing the activation in each voxel for studies containing the term (N = 182) compared with what one would expect if activation were uniformly distributed throughout the gray matter. **B.** Same format as A for a whole-brain FDR-corrected (*p* = 0.01) association-test meta-analysis z-score map of the “reasoning” term. This map was generated from a χ^2^ test comparing the proportion of studies demonstrating activation in each voxel for studies containing the term (N = 182) of interest compared with all other studies in the Neurosynth database (N > 14,000). Since the association test is more stringent than the uniformity test in A, it is unsurprising that there is less overlap between the left pimfs-v MPM and the clusters identified by the meta-analysis. Given the variability of the pimfs-v, future studies will determine if pimfs-v presence and functional regions in aLPFC related to reasoning are related to one another or not in individual participants.

**Supplementary References**

Yarkoni T, Poldrack RA, Nichols TE, et al (2011) Large-scale automated synthesis of human functional neuroimaging data. Nat Methods 8:665–670. https://doi.org/[10.1038/nmeth.1635](http://dx.doi.org/10.1038/nmeth.1635)
